# *SDHB* deficiency silences *ABCB1* expression and confers therapeutic vulnerability across multiple cancer types

**DOI:** 10.64898/2026.10.29.761968

**Authors:** Qianjin Guo, Anzel Zhen, Brooke E. Howitt, Justin P. Annes

## Abstract

Succinate dehydrogenase B (*SDHB*)-deficient cancers, including pheochromocytoma and paraganglioma (PPGL), renal cell carcinoma (RCC), and gastrointestinal stromal tumor (GIST), exhibit aggressive clinical behavior with limited treatment options. To identify novel therapeutic vulnerabilities, we performed high-throughput drug screening using co-cultured human *SDHB*-deficient and *SDHB*-reconstituted UOK269 human renal cell carcinoma cells with dual fluorescence labeling. Several compounds displaying preferential cytotoxicity toward *SDHB*-deficient cells were known substrates of the ATP-binding cassette B1 (ABCB1) transporter, also known as Multidrug Resistance Protein 1 (MDR1). Accordingly, *SDHB*-deficient cells demonstrated markedly reduced ABCB1 expression, a phenotype confirmed across human *SDHB*-deficient PPGL, RCC, and GIST samples, but absent in *IDH*- or *FH*-deficient tumors. Mechanistic investigation revealed that KDM4A histone demethylase is downregulated in *SDHB*-deficient tumors, and KDM4A loss (or inhibition) is sufficient to reduce ABCB1 expression, globally increase H3K9me3 levels, and sensitize cells to cytotoxic ABCB1 substrates. TCGA analysis confirmed correlated KDM4A-ABCB1 expression in *SDHB*-deficient cancers, with ABCB1 levels predicting drug sensitivity across cancer cell lines. Clinical analysis demonstrated that *SDHB*-mutant PPGL patients treated with vincristine-containing chemotherapy exhibited significantly longer progression-free survival compared to non-*SDHB* patients. These findings establish a novel metabolic-epigenetic pathway linking *SDHB* deficiency to ABCB1 silencing and provide rational basis for repurposing vincristine, and related ABCB1 substrate vinca alkaloids, across *SDHB*-deficient cancer types; offering an alternative precision treatment approach for these challenging malignancies.

## Introduction

Inherited mutations in succinate dehydrogenase genes (SDHx) cause the hereditary pheochromocytoma and paraganglioma (hPPGL) syndrome, which includes pheochromocytoma and paraganglioma (PPGL), renal cell carcinoma (RCC), and gastrointestinal stromal tumor (GIST). The mitochondrial SDH complex functions at the intersection of the tricarboxylic acid (TCA) cycle and electron transport chain (Complex II); catalyzing the conversion of succinate to fumarate while generating FADH₂ for oxidative phosphorylation (1). Loss of *SDHB* expression, or any SDH complex or assembly component, leads to succinate accumulation; an oncometabolite similar to other TCA intermediates like fumarate and 2-hydroxyglutarate (2-HG) (2–4), proposed to drive tumorigenesis *via* competitive inhibition of α-ketoglutarate–dependent dioxygenases, including prolyl-hydroxylases (PHD), JMJD2 family histone demethylases, and Ten-eleven translocation (TET) DNA demethylases (4–7). The tumorigenic consequences of oncometabolite accumulation are hypoxia-inducible factor (HIF) stabilization, genome wide epigenetic remodeling, and impaired DNA repair (2, 5, 8). Indeed, *SDHB*-deficient cancers (PPGL, RCC and GIST) (9–11), share cardinal molecular features of 1p loss (12, 13), succinate accumulation (14, 15), genome hypermethylation (5, 16, 17), and pseudohypoxic transcriptional signatures, despite distinct cellular origins (12). Raising the possibility of shared therapeutic susceptibilities across these distinct cancers.

*SDHB*-deficient tumors are particularly aggressive with frequent metastases and high mortality, making *SDHB* the most clinically significant SDHx gene (18–22). Current treatments for metastatic SDHB-PPGL include surgery, and systemic chemotherapeutics (e.g., cyclophosphamide, vincristine, dacarbazine), tyrosine kinase inhibitors (sunitinib, Cabozantinib, Axitinib, pazpanib), radiopharmaceutical medications (131I-mIBG), peptide receptor radionuclide therapy (PRRT, 177Lu-DOTATTE), and the recently approved HIF-2α inhibitor belzutifan (23–25). Unfortunately, these strategies typically demonstrate variable and limited efficacy (23, 26–28). Metastatic *SDHB*-RCC represents a subtype with poorly defined treatment approaches. While primary tumors are managed surgically, advanced disease is treated with conventional RCC therapies, including VEGF inhibitors and immune checkpoint inhibitors, though *SDHB*-RCC-specific efficacy data are limited (29–31). Epigenetic therapy with guadecitabine demonstrated biologic activity but not objective responses (32). Similarly, *SDHB*-GIST typically occurs in young patients and responds poorly to standard therapies. These tumors lack *KIT*/*PDGFRA* mutations, making them resistant to imatinib and related agents (33). Although recent studies suggest emerging activity of regorafenab and FGFR inhibition (34, 35), alternative agents including sunitinib, linsitinib, and guadecitabine show only modest efficacy (32, 36–38), Therefore, discovering novel therapeutic avenues for metastatic *SDHB*-cancers remains an unmet need.

In this study, we employed unbiased drug screening to identify therapeutic vulnerabilities in *SDHB*-deficient cancers. These experiments identified preferential sensitivity to chemotherapeutic compounds which are ATP-binding cassette transporter B1 (ABCB1/P-glycoprotein) substrates. Mechanistically, we found that *SDHB* deficiency silenced ABCB1 transporter expression, which promoted cytotoxic drug accumulation, through a novel mechanism involving Lysine Demethylase 4A (KDM4A) downregulation and, consequently, H3K9me3 accumulation. These findings reveal a targetable vulnerability of *SDHB*-deficient cancers with immediate clinical implications for precision therapy.

## Results

To identify agents with selective cytotoxicity towards *SDHB*-deficient cells we performed a high-throughput drug screen using co-cultured H2B-GFP-labeled *SDHB*-deficient UOK269 RCC cells (*SDHB*-KO) and isogenic H2B-mCherry-labeled *SDHB*-reconstituted control cells (WT) (Fig.1A) (39). Validation experiments confirmed that generation of the fluorescently labeled cell lines did not alter drug sensitivity, as demonstrated by comparable sensitivity of the parental and labeled cell lines to the chemotherapeutic compound Ym155; a drug we previously identified as having selective cytotoxicity towards SDH-deficient cells (Fig.1B) (40). For drug screening, we interrogated the arrayed NIH Clinical Collection library of 727 compounds at four concentrations (10, 100, 1000, 10,000 nM) in duplicate (Fig.1C). Following treatment (72 h), the relative viability of GFP-*SDHB*-KO and mCherry-WT cells was determined by automated high-content imaging. Multiple compounds demonstrated preferential cytotoxicity toward *SDHB*-KO cells (Fig.1D; Table S1). Top-ranked hits included actinomycin D, vinorelbine, vincristine, and chlorpromazine, which showed selectivity comparable to the positive control, Ym155. Subsequent validation of hit compounds with the parental *SDHB*-KO and WT cell lines confirmed selective cytotoxicity towards *SDHB*-KO cells for vinorelbine and vincristine (Fig.1E and Fig.S1). Notably, vinorelbine and vincristine are established substrates for the ATP-binding cassette transporter B1 (ABCB1/P-glycoprotein); a critical mediator of multidrug resistance that effluxes diverse chemotherapeutic agents including several anthracyclines, taxanes, and indole alkaloids (41–43). Mechanistically, ABCB1 reduces intracellular drug concentrations and contributes to treatment failure (44–47).

**Figure 1.**
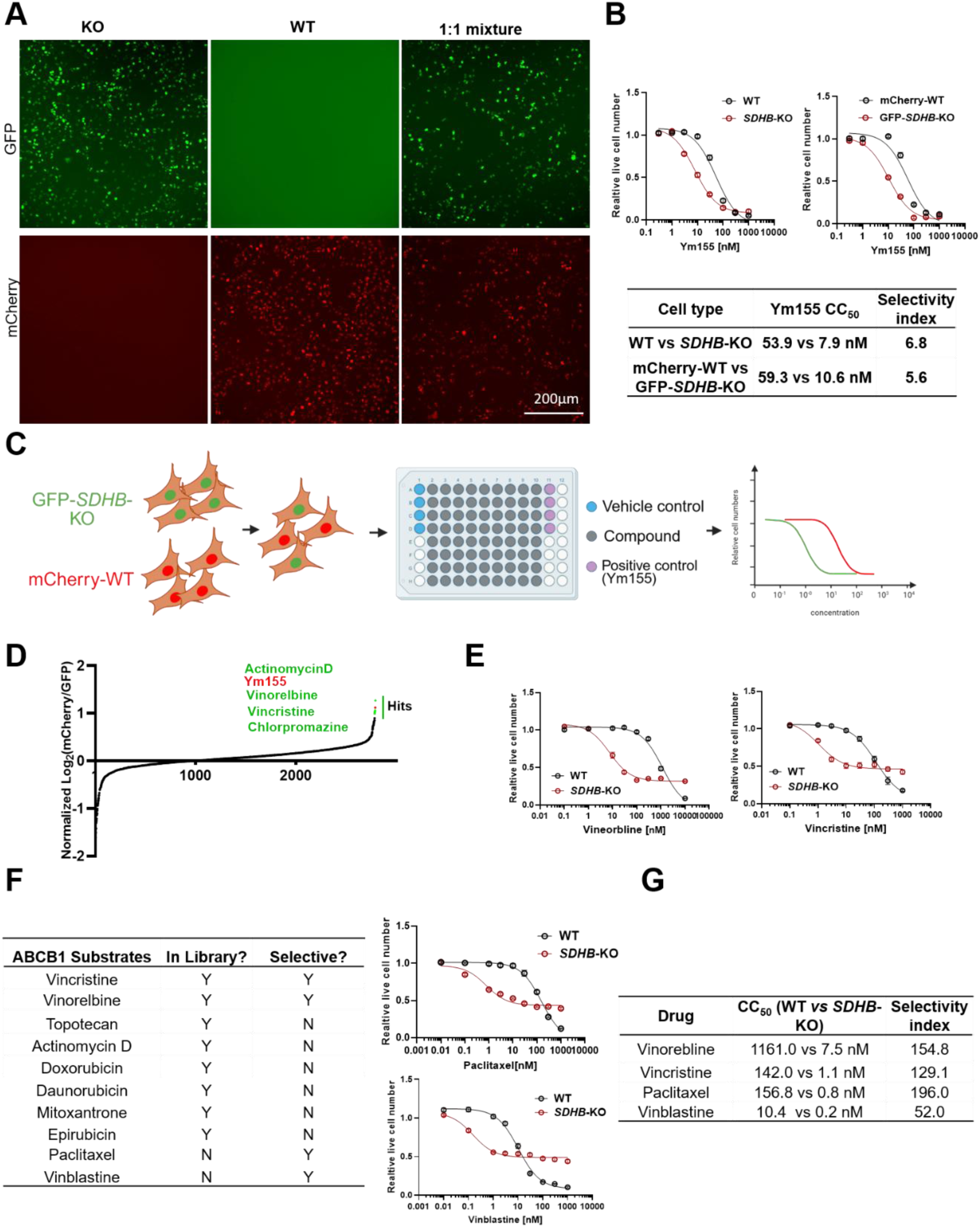
ABCB1 substrate drugs demonstrated selective toxicity in *SDHB*-deficient cells. **(A)** Representative images of UOK269 *SDHB*-deficient cells (GFP-*SDHB*-KO, green) and SDHB-reconstituted cells (mCherry-WT, red) cultured alone or co-cultured at a 1:1 ratio; scale bar = 200 μm. **(B)** Ym155 dose-response curves comparing labeled cells (GFP-*SDHB*-KO and mCherry-WT, right) with parental unlabeled cells (left). Table shows comparable CC₅₀ and selectivity values. **(C)** Schematic showing dual-labeled cell co-culture system screened against ∼800 compounds from NIH Clinical Collection library. **(D)** Drug screen results showed compounds with preferential cytotoxicity toward *SDHB*-deficient cells. Y-axis represents normalized selectivity score; X-axis shows individual compounds ranked by selectivity. Top hits (green), including actinomycin D, vinorelbine, vincristine, and chlorpromazine are highlighted. Ym155 (red) was the positive control compound. **(E)** Dose-response curves for vincristine (left) and vinorelbine (right) using the parental cell lines. **(F)** Table categorizing tested compounds by ABCB1 substrate status and selective cytotoxicity, and dose-response curves for paclitaxel and vinblastine in the parental cell lines. **(G)** Summary table of CC₅₀ values for validated compounds, demonstrating 50-200-fold greater potency in *SDHB*-deficient cells.

To determine whether *SDHB*-deficient cells exhibit broad sensitivity to ABCB1 substrates, we tested additional compounds including paclitaxel and vinblastine, and both demonstrated preferential cytotoxicity toward *SDHB*-KO cells (Fig.1F). However, not all ABCB1 substrates showed selective toxicity. Topoisomerase inhibitors such as topotecan, doxorubicin, and actinomycin D exhibited minimal differential effects. The selective vulnerability was primarily observed with microtubule-targeted agents, including vinca alkaloids (vincristine, vinorelbine, vinblastine) and taxanes (paclitaxel), which showed 50-200-fold greater potency in *SDHB*-deficient cells (Fig.1G).

To determine the molecular basis for ABCB1 substrate sensitivity in *SDHB*-KO cells, we examined ABCB1 transporter expression and function. *SDHB*-KO cells showed markedly reduced ABCB1 mRNA and protein levels compared to WT cells (Fig. 2A). Functional analysis using calcein acetoxymethyl (calcein AM) retention assays, a known ABCB1 substrate (48, 49), confirmed that reduced ABCB1 expression translated into impaired drug efflux activity (Fig. 2B, Fig. S2A). *SDHB*-KO cells showed significantly increased substrate accumulation compared to WT cells, and this difference was eliminated by treatment with the ABCB1 inhibitor elacridar; confirming the specificity of the transporter assay. To further verify that ABCB1 loss underlies drug hypersensitivity, we showed that elacridar treatment sensitized WT to vincristine to levels comparable to *SDHB*-KO cells (Fig. 2C). Hence, reduced ABCB1 expression mechanistically explained the preferential sensitivity of *SDHB*-KO cells to vincristine.

**Figure 2.**
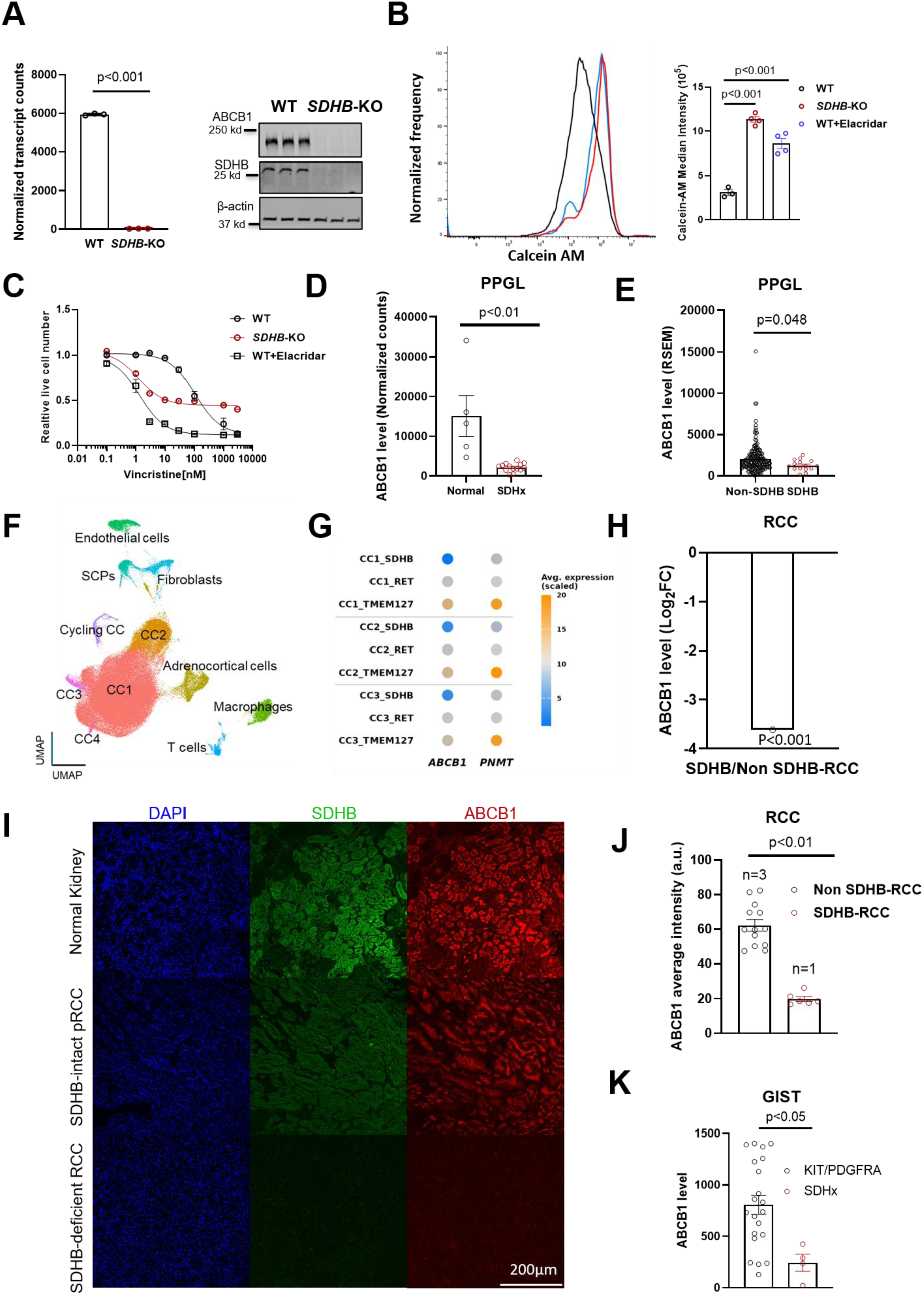
*SDHB* deficiency correlates with ABCB1 downregulation across multiple cancer types. **(A)** ABCB1 mRNA (left) and protein (right) expression in UOK269 cells comparing *SDHB*-deficiency (*SDHB*-KO) and SDHB-reconstituted (WT) cells. β-actin was loaded as control. ***p<0.001 by unpaired t-test. **(B)** Representative FACS data (left) and quantification analysis showed calcein AM accumulation in WT and *SDHB*-KO cells ± elacridar treatment (500 nM). **(C)** Vincristine dose-response curves in WT and *SDHB*-KO cells ± elacridar treatment (500 nM). **p<0.001 by unpaired t-test with Bonferroni correction. **(D)** ABCB1 mRNA expression in *SDHB*-mutant PPGL versus normal adrenal tissue from TCGA and GTEx databases. **p<0.001 by Mann-Whitney U test. **(E)** ABCB1 expression in *SDHB*-mutant PPGL (n=15) versus other PPGL genotypes (n=158) from TCGA. p=0.048 by Mann-Whitney U test. **(F)** UMAP projection of combined dataset (11 *SDHB*-mutant, 3 *TMEM127*-mutant, 2 *RET*-mutant tumors). **(G)** Dot plot of ABCB1 and PNMT expression in chromaffin cell populations (CC1-3) across genotypes. **(H)** ABCB1 expression in *SDHB*-deficient RCC versus other RCC subtypes. **p<0.001 by Mann-Whitney U test. **(I)** Immunofluorescence staining for ABCB1 in RCC samples. Representative images showing normal kidney, *SDHB*-intact RCC, and *SDHB*-deficient RCC, scale bar = 200 μm. **(J)** Quantification of the ABCB1 intensity from Figure 2I. **p<0.01 by Mann-Whitney U test. **(K)** ABCB1 expression in *SDHB*-deficient GIST versus other GIST subtypes from microarray analysis. **p<0.05 by Mann-Whitney U test.

Analysis of human *SDHB*-deficient tumors validated the translational relevance of these findings across human *SDHB*-deficient cancers. Human *SDHB*-PPGL showed significantly lower ABCB1 expression compared to normal adrenal tissue (Fig. 2D) and marginally lower expression compared to other PPGL genotypes (p=0.048) (Fig. 2E). To address potential cellular heterogeneity, we analyzed published single-cell RNA sequencing data of PPGL samples, combining 11 *SDHB*-mutant, 3 *TMEM127*-mutant, and 2 *RET*-mutant tumors (50, 51). UMAP clustering revealed distinct cellular populations, with cluster-specific marker genes confirming cell type identities (Fig. 2F, Fig. S2B). Chromaffin cell clusters (CC1-3) were identified by expression of tyrosine hydroxylase (TH) and phenylethanolamine N-methyltransferase (PNMT) (Fig. S2C-D). *SDHB*-mutant chromaffin cells showed consistently lower ABCB1 expression across clusters compared to *TMEM127*-mutant and *RET*-mutant samples (Fig. 2G). PNMT expression served as a positive control, as *SDHB*-mutant PPGLs characteristically exhibit a noradrenergic phenotype with low PNMT expression compared to the adrenergic phenotype of kinase signaling subtype PPGLs.

Extended analysis across *SDHB*-deficient cancer types confirmed the generality of ABCB1 downregulation. *SDHB*-deficient renal cell carcinoma showed significantly reduced ABCB1 expression compared to other RCC subtypes (Fig. 2H). Immunofluorescence analysis of RCC samples demonstrated ABCB*1* reduction at the protein level. When compared to normal kidney and *SDHB*-intact papillary RCC, *SDHB*-mutant RCC showed minimal ABCB1 staining (Fig. 2I and J). Microarray and RNAseq analysis of GIST samples similarly revealed lower ABCB1 expression in *SDHB*-deficient tumors compared to other genotypes (Fig. 2K and Fig.S2E). These results reveal ABCB1 downregulation as a consistent feature of *SDHB*-deficient cancers.

To determine whether ABCB1 downregulation is specific to *SDHB*-deficient cancers or a general feature of oncometabolite-related cancers, we analyzed ABCB1 expression in tumors with mutations in other TCA enzymes. Neither *IDH*-mutant glioblastomas nor *FH*-mutant renal cell carcinomas showed altered ABCB1 expression compared to their respective wild-type controls (Fig. 3A), demonstrating that *ABCB1* downregulation is specific to *SDHB* deficiency rather than a general consequence of metabolic reprogramming.

**Figure 3.**
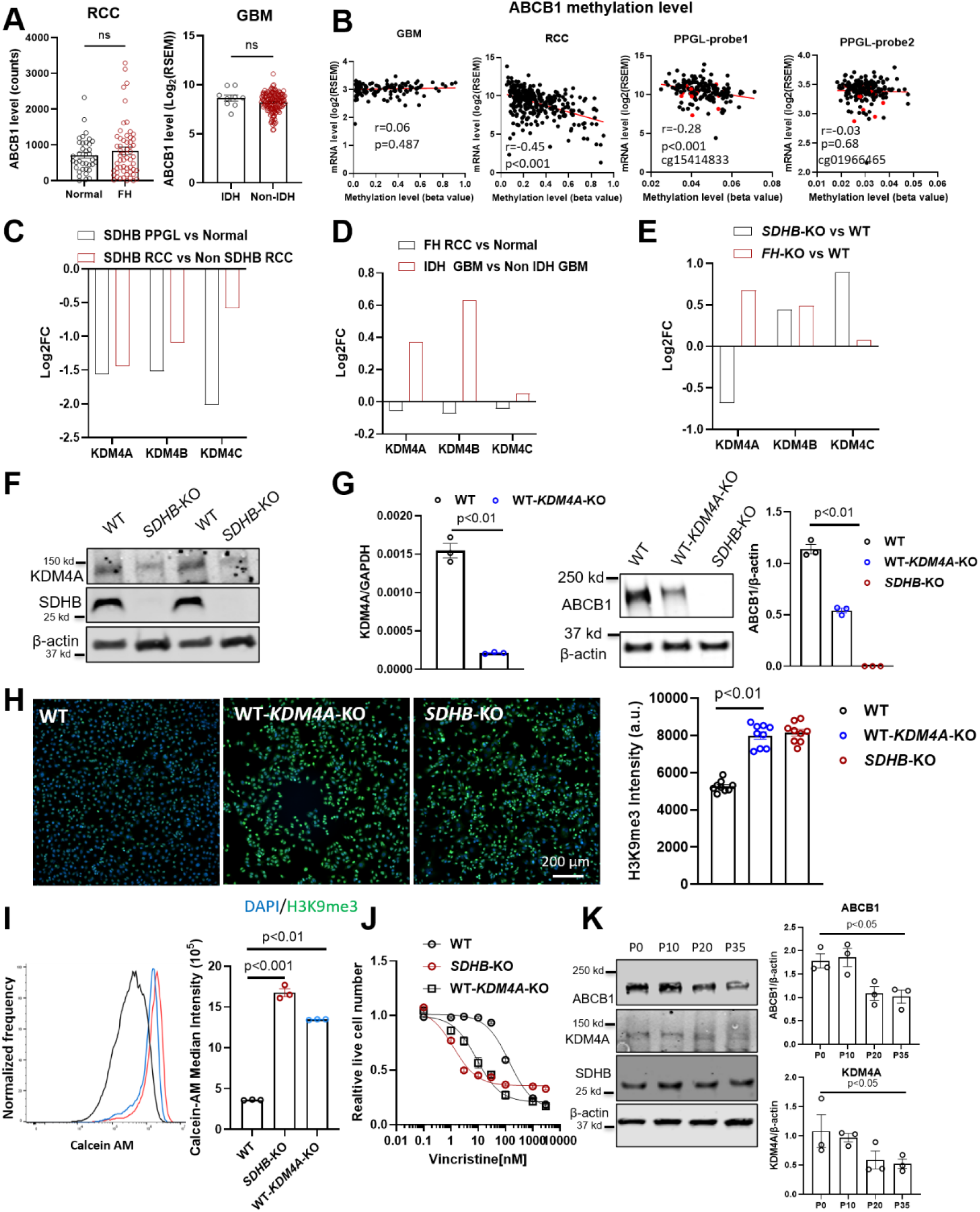
KDM4A downregulation drives ABCB1 silencing. **(A)** ABCB1 levels in *FH*-deficient RCC versus normal kidneys and *IDH*-mutant glioblastoma (GBM) versus other glioblastoma types, ns = not significant by Mann-Whitney U test. **(B)** Correlation between ABCB1 expression and promoter methylation in GBM, RCC, and PPGL (TCGA datasets). *SDHB*-mutant samples highlighted as red. Spearman correlation coefficients and p-values are indicated. **(C-D)** KDM4 family expression analysis. **(C)** KDM4A, KDM4B, and KDM4C expression were compared between *SDHB*-deficient and control samples (RCC and PPGL). **(D)** KDM4 family expression was analyzed in *FH*-deficient RCC and *IDH*-mutant glioblastoma. Box plots show log2 fold change. Statistical analysis was performed using Mann-Whitney U test. **(E)** KDM4 family expression in UOK269 (*SDHB*-KO) versus UOK262 (*FH*-KO) cell lines. **(F)** Western blot validation of KDM4A protein levels in *SDHB*-KO versus WT cells. β-actin serves as loading control. **(G)** KDM4A mRNA and ABCB1 protein expression following KDM4A knockout in *SDHB*-intact cells. **p<0.01 by unpaired t-test. **(H)** Global H3K9me3 levels in KDM4A-knockout cells. Immunofluorescence analysis showing increased H3K9me3 staining intensity in KDM4A-knockout cells compared to controls. Representative images and quantification from 3 independent experiments per condition, scale bar = 200 μm. **p<0.01 by unpaired t-test. **(I-J)** Functional consequences of KDM4A loss. **(I)** Calcein AM retention assay showing increased substrate accumulation in KDM4A-knockout cells. **(J)** Vincristine sensitivity comparing control and KDM4A-knockout cells. **(K)** Western blot analysis of KDM4A and ABCB1 expression in *SDHB*-KO cells treated with 3-NPA over passages (P0, P10, P20 and P35). Quantification shows progressive downregulation of both proteins. *p<0.05 by one-way ANOVA.

Although ABCB1 expression is reported to be regulated by DNA copy number changes, DNA methylation, and chromatin modifications, the regulatory mechanisms remain incompletely understood (52–57). Given that *ABCB1* promoter hypermethylation is a well-established mechanism of transcriptional silencing, we analyzed DNA methylation patterns across cancer types from TCGA. The correlation between ABCB1 expression and promoter methylation showed marked heterogeneity across cancer types (Fig. 3B). Interestingly, comparison of *SDHB*-mutant and *SDHB*-intact PPGL revealed no difference in *ABCB1* promoter methylation levels, and similarly, *SDHB*-KO and WT cells showed comparable ABCB1 methylation patterns (Fig. S3A), indicating that ABCB1 silencing in *SDHB*-deficient tumors is primarily DNA methylation independent.

To investigate alternative epigenetic mechanisms, we examined the KDM4 family of histone demethylases, which remove repressive H3K9me3 marks from gene promoters. Analysis of human tumor samples revealed that *KDM4A*, *KDM4B*, and *KDM4C* expression were all significantly downregulated in *SDHB*-mutant RCC and PPGL compared to controls (Fig. 3C). Importantly, this pattern was specific to *SDHB* deficiency, as *FH*-mutant RCC and *IDH*-mutant glioblastoma showed no reduction in KDM4 family expression (Fig. 3D), mirroring the specificity observed for *ABCB1*.

Cell line validation confirmed these findings, with *FH*-deficient UOK262 (*FH*-KO) cells showing minor change in *ABCB1* expression compared to controls (Fig. S3B), while *SDHB*-deficient UOK269 (*SDHB*-KO) cells demonstrated reduced KDM4A but not KDM4B or KDM4C expression (Fig. 3E-F). To verify causality, we generated *KDM4A* knockout cells in the *SDHB*-intact background and observed significant reduction in ABCB1 expression (Fig. 3G), accompanied by increased global H3K9me3 levels (Fig. 3H). Functionally, KDM4A loss recapitulated the *SDHB*-deficient phenotype, with enhanced calcein AM retention and increased vincristine sensitivity (Fig. 3I-J). Pharmacological validation using the KDM4 inhibitor SD49-7 provided additional mechanistic support. While short-term treatment showed no effect on vincristine sensitivity (Fig. S3C), one-week treatment produced minor differences in calcein AM retention and vincristine sensitivity (Fig. S3C-D). Extended two-week treatment resulted in elevated H3K9me3 levels, reduced ABCB1 expression, and significantly enhanced vincristine sensitivity (Fig. S3E-G), confirming that KDM4 inhibition phenocopies genetic KDM4A loss and supporting the role of this pathway in ABCB1 regulation.

To investigate whether sustained succinate accumulation recapitulates the *SDHB*-deficient phenotype, we treated cells with the SDH inhibitor 3-nitropropionic acid (3-NPA) over extended passages. While acute 3-NPA treatment (72 hours) failed to reduce ABCB1 expression (Fig. S3I), chronic treatment over 20 passages resulted in progressive downregulation of both KDM4A and ABCB1 (Fig. 3K). Similarly, temporal analysis of SDHB restoration revealed that short-term SDHB re-expression (48 hours) was insufficient to restore ABCB1 levels, whereas sustained SDHB restoration over one month significantly increased ABCB1 expression (Fig. S3J-K). These findings demonstrate that the SDHB-KDM4A-ABCB1 regulatory axis requires sustained metabolic reprogramming rather than acute perturbation.

To validate the clinical relevance of our mechanistic findings, we analyzed the relationship between KDM4A and ABCB1 expression across patient tumor samples from multiple *SDHB*-mutant cancer types. In PPGL tumors (TCGA cohort), KDM4A and ABCB1 expression were directly correlated (r=0.16, p=0.03, n=173), and this correlation was stronger (r value) within *SDHB*-mutant samples compared to the overall cohort; though significant, statistical power was limited by sample number (Fig. 4A). This correlation pattern was similarly observed in GIST samples where KDM4A and ABCB1 expression were also directly correlated (r=0.36, p=0.05, n=30) (Fig. 4B). The consistency of this biomarker relationship across multiple tumor types supports our mechanistic model linking KDM4A loss to ABCB1 downregulation.

**Figure 4.**
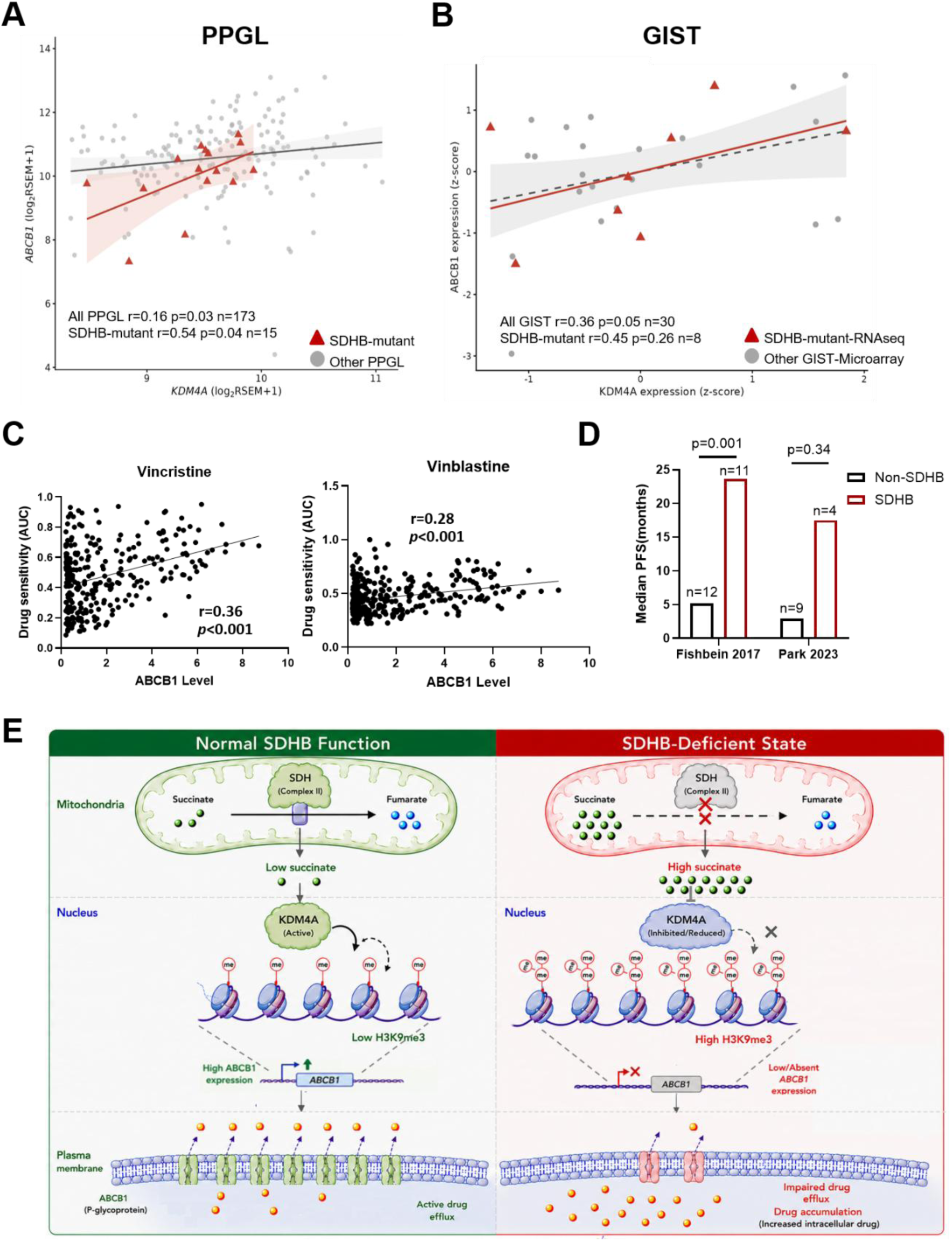
Clinical analysis of ABCB1/KDM4A biomarker signature across *SDHB*-deficient cancer types. **(A)** Correlation between KDM4A and ABCB1 expression in PPGL patient samples from TCGA (n=173). Each dot represents an individual tumor; red triangles indicate *SDHB*-mutant tumors, gray dots represent other PPGL genotypes. Linear regression line with 95% confidence interval shown. Spearman correlation coefficient and p-value indicated for all samples and *SDHB*-mutant subset. **(B)** KDM4A-ABCB1 correlation in GIST patient samples (n=30). *SDHB*-deficient tumors (red triangles) versus other GIST genotypes (gray dots). Linear regression with 95% confidence interval shown. Spearman correlation coefficient and p-value indicated for all samples and *SDHB*-deficient subset. **(C)** Scatter plots showing correlations between ABCB1 expression and drug sensitivity (AUC) for vincristine (left) and vinblastine (right). Each dot represents an individual cell line from Depmap. Lower AUC values indicate greater drug sensitivity. Pearson correlation coefficients and p-values indicated. **(D)** Progression-free survival (PFS) analysis in PPGL patients treated with CVD chemotherapy. Median PFS was compared between *SDHB*-mutant and non-SDHB PPGL patients in two independent cohorts: Fishbein 2017 (*SDHB* n=11, non-*SDHB* n=12) and Park 2023 (*SDHB* n=4, non-*SDHB* n=9). **(E)** Working model illustrating the SDHB–KDM4A–ABCB1 axis. *SDHB* deficiency causes succinate accumulation (green balls) and reduced KDM4A expression and activity, resulting in increased H3K9me3, reduced ABCB1 expression, impaired drug efflux (orange balls), and enhanced sensitivity to ABCB1 substrate drugs.

To functionally validate ABCB1 as a predictive biomarker for drug sensitivity, we analyzed the relationship between ABCB1 expression and microtubule agent sensitivity across diverse cancer cell lines. ABCB1 expression showed significant correlations with sensitivity to both vincristine (r=0.36, p<0.001) and vinblastine (r=0.28, p<0.001), confirming that ABCB1 levels serve as a robust predictor of response to these agents across cancer cell type (Fig. 4C). To verify the specificity of this relationship, we examined correlations between ABCB1 expression and sensitivity to standard RCC therapies that are not ABCB1 substrates. Lenvatinib, sunitinib, and everolimus showed no correlation with ABCB1 expression levels (Fig. S4), confirming the predictive utility of ABCB1 is specific to its substrates rather than representing a general chemosensitivity marker.

To assess the clinical relevance of ABCB1 downregulation in *SDHB*-deficient tumors, we retrieved progression-free survival (PFS) data from PPGL patients treated with CVD chemotherapy. *SDHB*-mutant PPGLs demonstrated significantly longer median PFS compared to non-*SDHB* tumors in the Fishbein 2017 cohort (23.67 vs 5.17 months, p=0.001, n=11 vs n=12) (27), while the Park 2023 cohort (58) showed a similar trend (17.50 vs 2.90 months, p=0.34, n=4 vs n=9) (Figure 4D). Although limited by the rarity of *SDHB*-mutant tumors (n=15), this analysis suggests potential prognostic utility of ABCB1 expression in this patient population.

## Discussion

Using unbiased high-throughput chemical screening we identified a previously unrecognized therapeutic vulnerability in *SDHB*-deficient cancers mediated by the selective downregulation of the multidrug resistance gene drug *ABCB1*. *SDHB* loss reduces ABCB1 expression, creating hypersensitivity to microtubule-targeting transporter substrates that is recapitulated by impaired KDM4A activity (Figure 4E). ABCB1 downregulation is conserved across multiple *SDHB*-deficient cancer types (PPGL, RCC, and GIST), providing an actionable therapeutic strategy for treating these challenging malignancies.

An intriguing finding of this study is that suppression of the ABCB1 was observed in *SDHB*-deficient tumors but not in *FH*- or *IDH*-deficient models. While *SDHB*, *IDH*, and *FH* mutations all cause oncometabolite accumulation, genome-wide DNA and histone hypermethylation (5, 59–61), only *SDHB* loss resulted in KDM4A downregulation and consequent ABCB1 silencing. Our data underscore the importance of individually conceptualizing the mechanisms of oncometabolite-related tumorigenesis, consistent with current evidence indicating that oncometabolites exert differential downstream target effects. This specificity likely reflects multiple mechanistic factors. First, despite shared global hypermethylation patterns, gene-specific methylation effects vary substantially between oncometabolites. Our analysis revealed no difference in ABCB1 promoter methylation between *SDHB*-mutant and other PPGL subtypes, while *IDH*-mutant glioblastomas showed ABCB1 hypomethylation compared to wild-type tumors (62). Although both *FH*- and *SDHB*-deficient RCC display global DNA hypermethylation, *SDHB* tumors exhibit a distinct methylation landscape from *FH* tumors (63). Second, recent studies have demonstrated that ABCB1 expression can be regulated through higher-order chromatin organization and enhancer rewiring independent of promoter DNA methylation (53). Our findings extend this concept by identifying that reduced KDM4A expression correlates with ABCB1 suppression in *SDHB*-deficient tumors. The correlation between reduced KDM4 expression, increased H3K9me3 levels, and ABCB1 suppression supports a histone-mediated regulatory mechanism as a contributing factor in *SDHB*-specific gene silencing.

Third, different oncometabolites exhibit distinct potencies against downstream chromatin-modifying targets. Biochemical studies have shown that succinate is a more potent KDM4A inhibitor than fumarate (6). Meanwhile, 2-hydroxyglutarate shows variable inhibition dynamics across different α-KG-dependent enzymes (61). The combination of gene-specific epigenetic targeting, histone-mediated regulation and differential biochemical potencies likely explains why *SDHB* deficiency uniquely suppresses ABCB1 expression.

We identify KDM4A inhibition as a key link between *SDHB* loss and *ABCB1* silencing. KDM4A demethylase removes repressive H3K9me3 marks, and its down-regulation in *SDHB*-deficient cells leads to enhanced H3K9me3 accumulation and transcriptional silencing of *ABCB1* expression. While we demonstrate that KDM4A loss is sufficient to recapitulate this *SDHB*-deficient phenotype, the mechanism by which SDHB regulates KDM4A expression remains unclear. Chronic 3-NPA treatment progressively downregulated KDM4A expression in *SDHB* wild-type cells, suggesting that sustained succinate elevation and/or KDM4A suppression is required to expert epigenetic reprogramming. Understanding the mechanistic details of how KDM4A activity is regulated will require further investigation.

Our findings have immediate clinical implications for *SDHB*-deficient cancer management. *SDHB*-mutant PPGLs demonstrated significantly longer progression-free survival with CVD treatment compared to non-*SDHB* PPGLs (23 vs 5 months, p=0.001) (27); our findings provide molecular rationale for this clinically important observation and suggest that *SDHB* mutation status combined with ABCB1 expression might identify patients most likely to benefit from vincristine-based therapy. Interestingly, the futility of treating RCC with chemotherapy-including taxanes-is well-established (64–66). However, based upon our data, we propose trialing use of vincristine (or vinblastine) for *SDHB*-deficient RCC and GIST; these cancer subtypes currently lack effective treatments and resist standard therapies. The substantial drug sensitivity differences observed (50-200-fold) suggest that prior treatment failures with these therapeutics may have masked potential efficacy in this rare subtype with greatly reduced ABCB1 expression (which could be confirmed immunohistochemically). We propose implementation of ABCB1/KDM4A biomarker assessment, following *SDHB* mutation testing to enable a precision medicine approach; directing patients with low ABCB1 expression towards specific chemotherapeutic treatment.

In conclusion, we identified and characterized a novel therapeutic vulnerability in *SDHB*-deficient cancers that creates immediate opportunities for precision medicine implementation. The SDHB-KDM4A-ABCB1 regulatory axis represents a paradigm for understanding how metabolic tumor suppressors influence drug responses through specific epigenetic mechanisms. Our findings provide molecular rationale for repurposing vincristine and related agents across multiple *SDHB*-deficient cancer types. Implementation of ABCB1/KDM4A biomarker testing could guide precision medicine approaches in these aggressive malignancies.

## Materials and Methods

### Cell Culture

UOK269 cell lines were provided by Dr. W Marston Linehan and maintained in DMEM (Cytiva, SH30243.01) with addition of 10 % fetal bovine serum (FBS,Cytiva,SH30396.03HI), 100 U/mL penicillin-streptomycin (Corning, 30-002-Cl) and 1X Non-Essential Amino Acid (NEAA) Solution (Cytiva,SH30238.01). The cell line was used within 20 passages.

### High-throughput drug screening

UOK269 (*SDHB*-KO) and UOK269-SDHB (WT) cells were stably transduced to express nuclear H2B-tagged fluorescent proteins, which have been widely utilized in high-throughput drug screenings (67, 68). NIH Clinical Collection library (727 compounds) was used for screening. Each well of a 96-well plate was seeded with a mixture of 5,000 H2B-GFP-KO cells and 5,000 H2B-mCherry-WT cells. Four concentrations of each compound were tested: 10 nM, 100 nM, 1,000 nM, and 10,000 nM, with duplicates for each concentration. Ym155 chemotherapy and DMSO were used as positive control and negative control, respectively. After 72 hours, cells were imaged by Operetta CLS High-Content Analysis System using the GFP and mCherry channels to assess the cell numbers. The ratio of mCherry positive numbers to GFP positive numbers was quantified for analysis.

### Immunofluorescence Staining

Cells were seeded in 96-well plates, treated as indicated, and fixed with 4% paraformaldehyde for 10 minutes at room temperature. Following permeabilization with 0.1% Triton X-100 for 10 minutes and blocking with 1% donkey serum for 1 hour, cells were incubated overnight at 4°C with primary antibodies against H3K9me3 (Active Motif, Cat #39062), SDHB (Abcam, Cat #ab14714), and ABCB1 (Proteintech, Cat #22336-1-AP). After washing with PBS, cells were incubated with Alexa Fluor 647-conjugated secondary antibodies for 1 hour at room temperature. Nuclei were counterstained with DAPI (1 μg/ml). Images were acquired using an Operetta CLS High-Content Imaging System and analyzed with Harmony 4.5 software (Revvity).

### Flow cytometry assessment of calcein AM retention

To assess ABCB1 efflux activity, 3 × 10⁵ cells were resuspended in PBS containing 10 nM freshly prepared calcein AM (Fisher Scientific, Cat #C3099) with either 500 nM elacridar (ABCB1 inhibitor) or vehicle control. Samples were incubated for 20 minutes at 37°C, then resuspended in prewarmed culture medium and incubated for an additional 10 minutes to ensure optimal dye retention. Calcein-AM accumulation was measured by flow cytometry using an Acea Novocyte Quanteon cytometer (Agilent). Data analysis was performed using FlowJo version 10.1 (BD Biosciences).

### Cell Viability Assay

Cells were seeded in 96-well plates (10,000 cells per well) and treated with serial dilutions of compounds for 72 hours. Cell viability was assessed by staining with DAPI and Ethidium Homodimer I (EthD-1, Biotium) for 30 minutes at room temperature. Images were acquired using an Operetta CLS High-Content Imaging System and analyzed with Harmony 4.5 software (Revvity). Viable cells were identified as DAPI-positive/EthD-1-negative and normalized to vehicle-treated controls. Dose-response curves and CC₅₀ values were calculated using GraphPad Prism software.

### Western Blotting (WB)

Cells were lysed in RIPA buffer (CST, Cat #9806S) containing protease and phosphatase inhibitors (Thermo Fisher, Cat #78440). Protein concentration was determined using a BCA assay (Pierce). Equal amounts of protein (20 μg) were separated by SDS-PAGE and transferred to PVDF membranes. Membranes were blocked with 1% BSA in Intercept® Blocking Buffer (LI-COR, Cat #927-60001) for 1 hour at room temperature and incubated overnight at 4°C with primary antibodies against SDHB (Abcam, Cat #ab14714), ABCB1 (Proteintech, Cat #22336-1-AP), KDM4A (CST, Cat #5328T), and β-actin (Sigma, Cat #A5316). Following three washes with TBST, membranes were incubated with IRDye-conjugated secondary antibodies (1:10,000) for 1 hour at room temperature. Blots were imaged using the Odyssey imaging system (LI-COR).

### CRISPR/Cas9-Mediated KDM4A Knockout

To generate KDM4A knockout cells, a single-guide RNA (sgRNA) targeting human KDM4A was cloned into the pMCB320 lentiviral vector. Lentiviral particles were produced by co-transfecting HEK293T cells with pMCB320-sgKDM4A, pRSV-Rev, pMDLg/pRRE, and pMD2.G plasmids using Lipofectamine 3000. Viral supernatants were harvested 48 hours post-transfection, filtered through a 0.45 μm filter, and used to transduce UOK269 SDHB-reconstituted cells (WT) in the presence of 8 μg/ml polybrene. After 48 hours, mCherry-positive cells were sorted by flow cytometry and expanded for downstream validation. KDM4A-gRNA-F 5’-TTGGAGTGAACTGCCTCCAAGAGCGTTTAAGAGC-3’; KDM4A-gRNA-R 5’-TTAGCTCTTAAACGCTCTTGGAGGCAGTTCACTCCAACAAG-3’

### Methylation Analysis

Genomic DNA was extracted from UOK269 cells using standard protocols. For methylation analysis, 100 ng of genomic DNA was subjected to sodium bisulfite conversion using the EZ DNA Methylation-Direct Kit (Zymo Research) according to manufacturer’s instructions. Methylation-specific PCR (MSP) primers, targeting the ABCB1 promoter region, cover key CpG sites within the functional promoter sequence as previously reported (69). PCR amplification was performed using bisulfite-converted DNA with primers specific for methylated and unmethylated sequences. The primer sequences were: methylated-specific forward (M-F): CGAGGAATTAGTATTTAGTTAATTCGGGTCGG, methylated-specific reverse (M-R): ACTCAACCCACGCCCCGACG, unmethylated-specific forward (U-F): TGAGGAATTAGTATTTAGTTAATTTGGGTTGG, unmethylated-specific reverse (U-R): ACTCAACCCACACCCCAACA. PCR products were analyzed by electrophoresis on 2% agarose gels and visualized using ethidium bromide staining. Methylation status was determined by the presence or absence of specific PCR products. Negative controls without DNA template were included in each experiment

### Public Dataset Analysis

Gene expression and clinical data were obtained from The Cancer Genome Atlas (TCGA) and other publicly available databases. For PPGL and glioblastoma (GBM) samples, RNA-seq data were downloaded from TCGA-PCPG and TCGA-GBM cohorts, respectively, with samples classified based on *SDHB* mutation status using previously published genomic annotations (13). GIST and RCC datasets were obtained from published studies (12, 70). Single-cell RNA sequencing data were processed using Seurat and integrated data from multiple studies (50, 71). RNA-seq data for *FH*-deficient RCC and UOK262 cell lines were obtained from Gene Expression Omnibus (GEO) datasets GSE273036 and GSE77538 (60, 72), respectively. DNA methylation data for PPGL, RCC, and GBM were obtained from TCGA. Beta values were used for correlation analysis between promoter methylation and gene expression levels.

### Statistical analysis

Results are expressed as mean ± SE. Statistical analysis was performed using Student’s t-test, Mann–Whitney U test, or one-way ANOVA with repeated measures. A value of *p* <0.05 was considered significant.

## Acknowledgments

This study was supported in part by a philanthropic gift from the Lau Family to advance research on SDHB-related disease, and by research grants from the PARADIFFERENCE foundation (JPA) and SDHB PheoPara Coalition (JPA). Additional grant support was received from the NIH NIDDK (JPA: U01DK136965, R01DK101530 and R01DK119955), the Neuroendocrine Tumor Research Foundation (QG), Allen and Naomi Schneider Summer Research Fund (AZ).

## Supplementary information

**Supplementary Figure S1.**
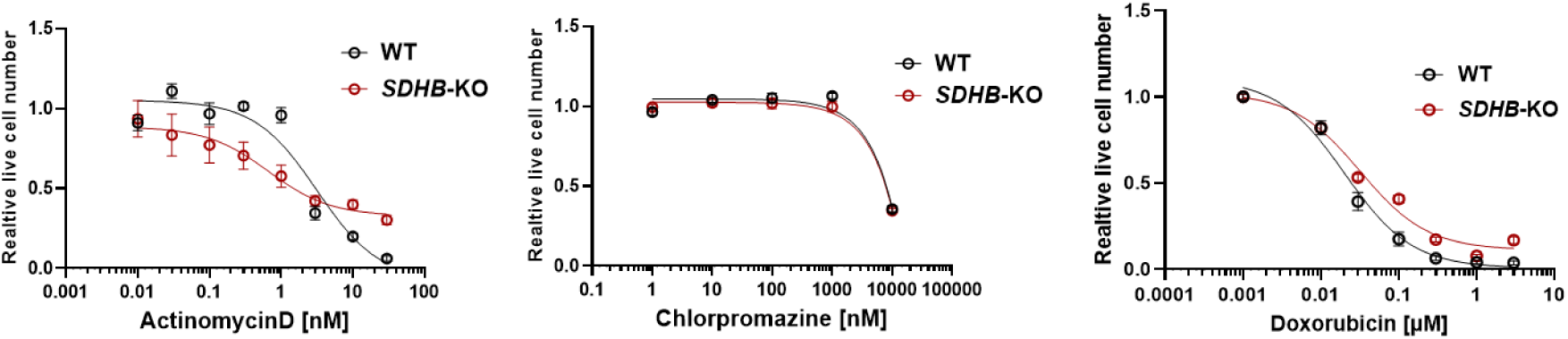
Dose-response curves showing relative cell viability after treatment with actinomycin D (left) and chlorpromazine (middle) and doxorubicin (right) in UOK269 *SDHB* wild-type (WT) and deficient (*SDHB*-KO) cells. Data represent mean ± SEM from three independent experiments.

**Table S1. Summary of the drug screening results.**

**Supplementary Figure S2.**
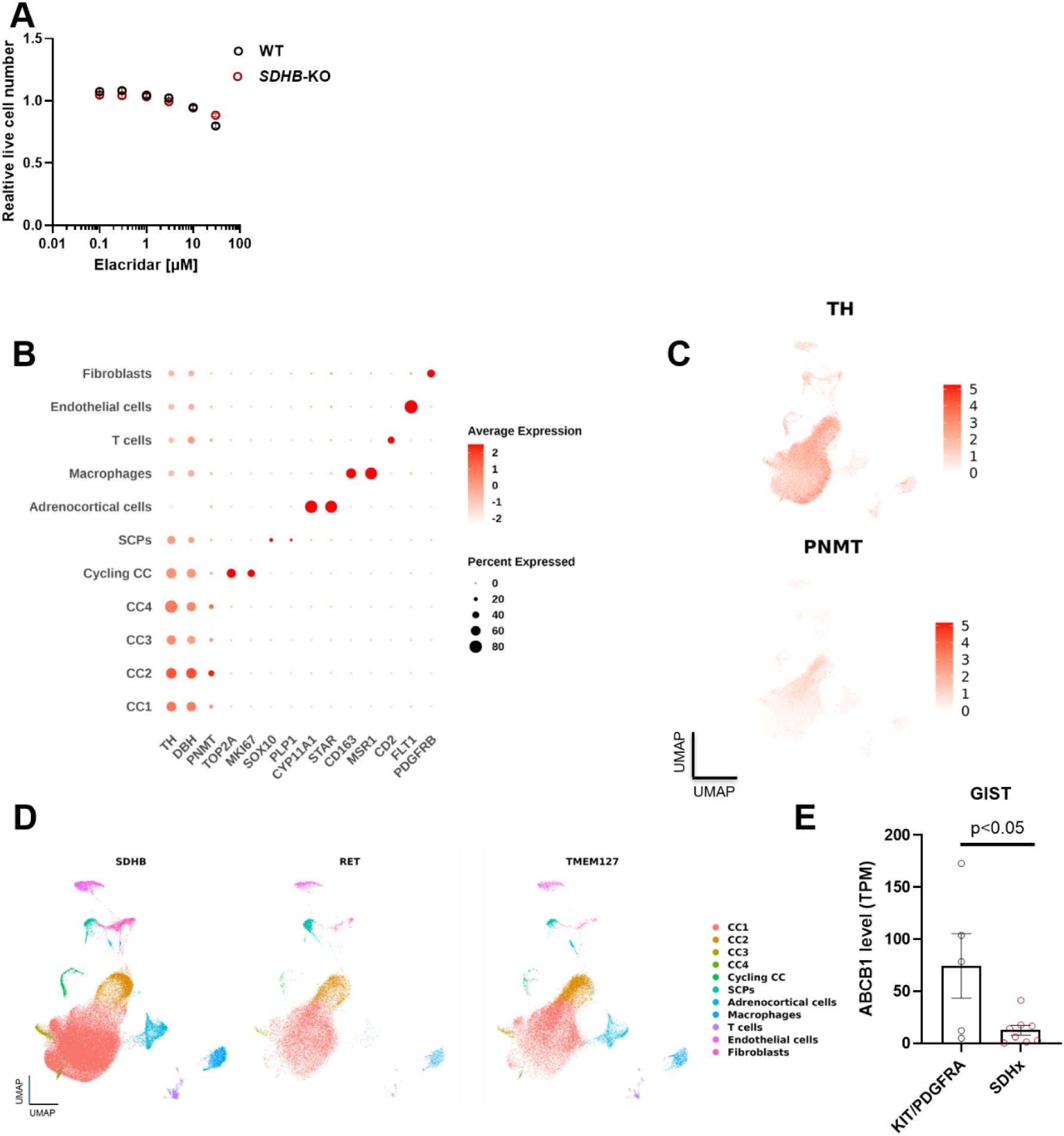
ABCB1 regulation and single-cell analysis of PPGL samples. **(A)** Dose-response analysis of elacridar cytotoxicity in WT and *SDHB*-KO cells. **(B)** Dot plot showing cluster-specific marker genes from single-cell RNA sequencing analysis of PPGL samples. Data from combined analysis of 11 *SDHB*-mutant, 3 *TMEM127*-mutant, and 2 *RET*-mutant PPGL samples. **(C)** UMAP projections showing expression of chromaffin cell markers tyrosine hydroxylase (TH, top) and phenylethanolamine N-methyltransferase (PNMT, bottom). **(D)** Genotype-stratified UMAP projections showing cellular composition by mutation type. Left: *SDHB*-mutant, Middle: *RET*-mutant, Right: *TMEM127*-mutant PPGL samples. **(E)** ABCB1 expression in *SDHB*-deficient GIST versus other GIST subtypes from mRNA-seq analysis. *p<0.05 by Mann-Whitney U test.

**Supplementary Figure S3.**
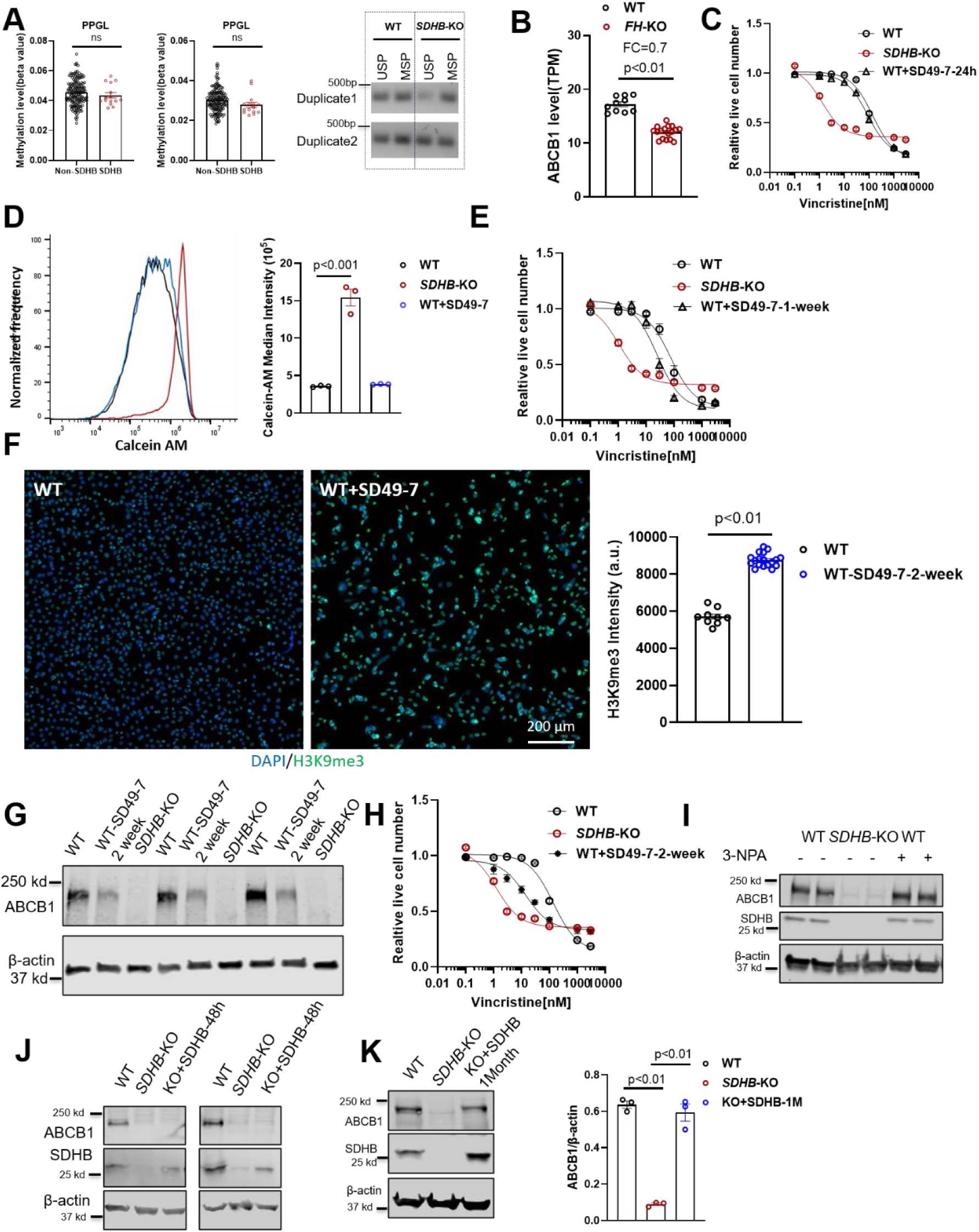
Pharmacological KDM4 inhibition phenocopies genetic KDM4A loss. **(A)** ABCB1 promoter methylation in *SDHB*-deficient versus non-SDHB samples. Methylation levels in PPGL patient tumors from TCGA (left two panels) and UOK269 cells (right panel). Individual dots represent tumor samples. No significant methylation differences were detected between genotypes. Statistical analysis by Mann-Whitney U test. **(B)** Comparison of ABCB1 levels between *FH*-deficient UOK262 (*FH*-KO) and WT cells. **p<0.01 by Mann-Whitney U test. **(C)** Vincristine sensitivity in WT cells treated with SD49-7 (KDM4 inhibitor) for 24 hours. **(D)** Calcein AM retention assay and **(E)** vincristine sensitivity in WT cells treated with SD49-7 for one week. **(F)** H3K9me3 levels following two-week SD49-7 treatment. Immunofluorescence analysis showing increased H3K9me3 in treated versus control cells, scale bar = 200 μm. **p<0.01 by Mann-Whitney U test. **(G)** ABCB1 protein levels after two-week SD49-7 treatment. β-actin serves as loading control. **(H)** Vincristine sensitivity following two-week SD49-7 treatment. Dose-response curves show enhanced sensitivity in inhibitor-treated cells. **(I)** Western blot analysis of ABCB1 in *SDHB*-KO cells treated with 3-NPA for 72 hours. **(J)** Western blot analysis of ABCB1 in *SDHB*-KO cells treated with short-term SDHB re-expression (48 hours). **(K)** Western blot analysis of ABCB1 in *SDHB*-KO cells with SDHB restoration over one month. quantification from three independent experiments. **p<0.01 by unpaired t-test with Bonferroni correction.

**Supplementary Figure S4.**
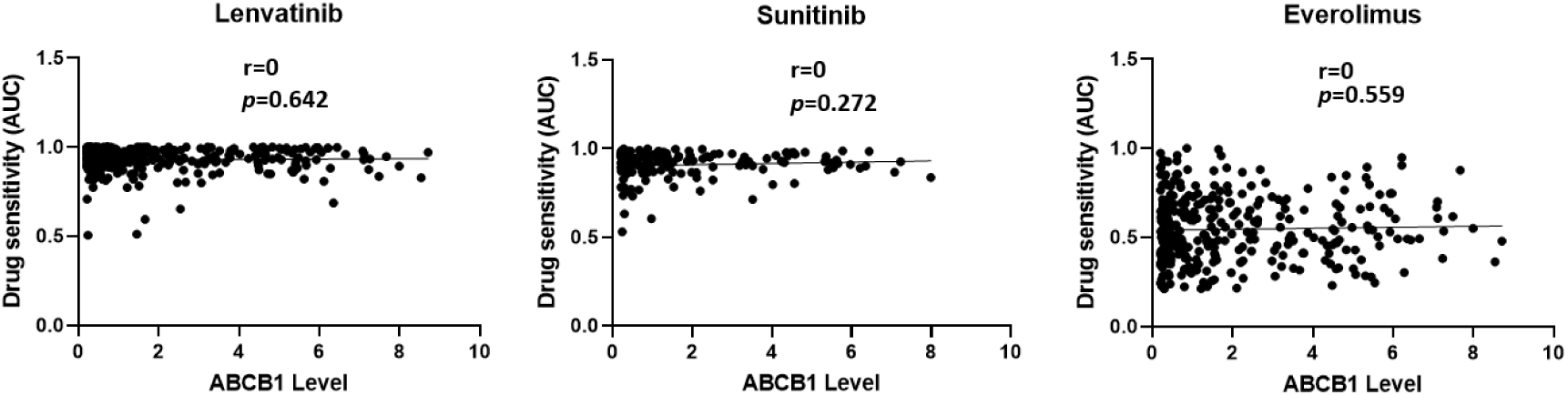
Scatter plots show correlations between ABCB1 expression and drug sensitivity (AUC) for Lenvatinib, Sunitinib and Everolimus. Each dot represents an individual cell line from Depmap. Lower AUC values indicate greater drug sensitivity. Pearson correlation coefficients and p-values indicated.

## References

1. B. A. Ackrell, M. K. Johnson, R. P. Gunsalus, G. Cecchini, "Structure and function of succinate dehydrogenase and fumarate reductase" in Chemistry and biochemistry of flavoenzymes. (CRC Press, 2019), pp. 229–297.

2. M. A. Selak et al., Succinate links TCA cycle dysfunction to oncogenesis by inhibiting HIF-alpha prolyl hydroxylase. Cancer Cell 7, 77–85 (2005).

3. L. Dang et al., Cancer-associated IDH1 mutations produce 2-hydroxyglutarate. Nature 462, 739–744 (2009).

4. N. Armstrong et al., SDHB knockout and succinate accumulation are insufficient for tumorigenesis but dual SDHB/NF1 loss yields SDHx-like pheochromocytomas. Cell Rep 38, 110453 (2022).

5. E. Letouze et al., SDH mutations establish a hypermethylator phenotype in paraganglioma. Cancer Cell 23, 739–752 (2013).

6. M. Xiao et al., Inhibition of alpha-KG-dependent histone and DNA demethylases by fumarate and succinate that are accumulated in mutations of FH and SDH tumor suppressors. Genes Dev 26, 1326–1338 (2012).

7. P. L. Sulkowski et al., Oncometabolites suppress DNA repair by disrupting local chromatin signalling. Nature 582, 586–591 (2020).

8. P. L. Sulkowski et al., Krebs-cycle-deficient hereditary cancer syndromes are defined by defects in homologous-recombination DNA repair. Nat Genet 50, 1086–1092 (2018).

9. K. A. Andrews et al., Tumour risks and genotype-phenotype correlations associated with germline variants in succinate dehydrogenase subunit genes SDHB, SDHC and SDHD. J Med Genet 55, 384–394 (2018).

10. S. Vanharanta et al., Early-onset renal cell carcinoma as a novel extraparaganglial component of SDHB-associated heritable paraganglioma. Am J Hum Genet 74, 153–159 (2004).

11. H. P. Neumann et al., Distinct clinical features of paraganglioma syndromes associated with SDHB and SDHD gene mutations. JAMA 292, 943–951 (2004).

12. J. B. Neves et al., Defining the origin, evolution, and immune composition of SDH-deficient renal cell carcinoma. iScience 25, 105389 (2022).

13. L. Fishbein et al., Comprehensive Molecular Characterization of Pheochromocytoma and Paraganglioma. Cancer Cell 31, 181–193 (2017).

14. S. Lee et al., Functional Characterization of SDHB Variants Clarifies Hereditary Pheochromocytoma and Paraganglioma Risk and Genotype-Phenotype Relationships. J Clin Invest 10.1172/JCI198165 (2025).

15. E. Kim et al., Utility of the succinate: Fumarate ratio for assessing SDH dysfunction in different tumor types. Mol Genet Metab Rep 10, 45–49 (2017).

16. W. A. Flavahan et al., Altered chromosomal topology drives oncogenic programs in SDH-deficient GISTs. Nature 575, 229–233 (2019).

17. J. K. Killian et al., Succinate dehydrogenase mutation underlies global epigenomic divergence in gastrointestinal stromal tumor. Cancer Discov 3, 648–657 (2013).

18. S. Hescot et al., Prognosis of Malignant Pheochromocytoma and Paraganglioma (MAPP-Prono Study): A European Network for the Study of Adrenal Tumors Retrospective Study. J Clin Endocrinol Metab 104, 2367–2374 (2019).

19. J. Crona et al., Genotype-phenotype correlations in pheochromocytoma and paraganglioma: a systematic review and individual patient meta-analysis. Endocr Relat Cancer 26, 539–550 (2019).

20. C. Horton et al., Universal Germline Panel Testing for Individuals With Pheochromocytoma and Paraganglioma Produces High Diagnostic Yield. J Clin Endocrinol Metab 107, e1917– e1923 (2022).

21. R. Jabarkheel et al., Metastatic Paraganglioma of the Spine With SDHB Mutation: Case Report and Review of the Literature. Int J Spine Surg 14, S37–S45 (2021).

22. C. J. Ricketts et al., Succinate dehydrogenase kidney cancer: an aggressive example of the Warburg effect in cancer. J Urol 188, 2063–2071 (2012).

23. D. Taieb et al., Management of phaeochromocytoma and paraganglioma in patients with germline SDHB pathogenic variants: an international expert Consensus statement. Nat Rev Endocrinol 20, 168–184 (2024).

24. C. Jimenez et al., Belzutifan for Advanced Pheochromocytoma or Paraganglioma. N Engl J Med 393, 2012–2022 (2025).

25. R. Garcia-Carbonero et al., Multidisciplinary practice guidelines for the diagnosis, genetic counseling and treatment of pheochromocytomas and paragangliomas. Clin Transl Oncol 23, 1995–2019 (2021).

26. H. Huang et al., Treatment of malignant pheochromocytoma/paraganglioma with cyclophosphamide, vincristine, and dacarbazine: recommendation from a 22-year follow-up of 18 patients. Cancer 113, 2020–2028 (2008).

27. L. Fishbein et al., SDHB mutation carriers with malignant pheochromocytoma respond better to CVD. Endocr Relat Cancer 24, L51–L55 (2017).

28. L. Fishbein et al., The North American Neuroendocrine Tumor Society Consensus Guidelines for Surveillance and Management of Metastatic and/or Unresectable Pheochromocytoma and Paraganglioma. Pancreas 50, 469–493 (2021).

29. T. H. Tsai, W. Y. Lee, Succinate Dehydrogenase-Deficient Renal Cell Carcinoma. Arch Pathol Lab Med 143, 643–647 (2019).

30. M. I. Carlo et al., Familial Kidney Cancer: Implications of New Syndromes and Molecular Insights. Eur Urol 76, 754–764 (2019).

31. R. R. Kotecha et al., A Phase 2 Trial of Talazoparib and Avelumab in Genomically Defined Metastatic Kidney Cancer. Eur Urol Oncol 7, 804–811 (2024).

32. J. A. Ligon et al., A Phase II Trial of Guadecitabine in Children and Adults with SDH-Deficient GIST, Pheochromocytoma, Paraganglioma, and HLRCC-Associated Renal Cell Carcinoma. Clin Cancer Res 29, 341–348 (2023).

33. A. Ibrahim, S. Chopra, Succinate Dehydrogenase-Deficient Gastrointestinal Stromal Tumors. Arch Pathol Lab Med 144, 655–660 (2020).

34. P. Merriam et al., Fibroblast growth factor receptor inhibition for succinate dehydrogenase-deficient gastrointestinal stromal tumors: a phase 2 trial. Nat Med 10.1038/s41591-026-04376-9 (2026).

35. J. Martin-Broto et al., REGISTRI: Regorafenib in first-line of KIT/PDGFRA wild type metastatic GIST: a collaborative Spanish (GEIS), Italian (ISG) and French Sarcoma Group (FSG) phase II trial. Mol Cancer 22, 127 (2023).

36. K. A. Janeway et al., Sunitinib treatment in pediatric patients with advanced GIST following failure of imatinib. Pediatr Blood Cancer 52, 767–771 (2009).

37. E. Ben-Ami et al., Long-term follow-up results of the multicenter phase II trial of regorafenib in patients with metastatic and/or unresectable GI stromal tumor after failure of standard tyrosine kinase inhibitor therapy. Ann Oncol 27, 1794–1799 (2016).

38. M. von Mehren et al., Linsitinib (OSI-906) for the Treatment of Adult and Pediatric Wild-Type Gastrointestinal Stromal Tumors, a SARC Phase II Study. Clin Cancer Res 26, 1837– 1845 (2020).

39. N. Saxena et al., SDHB-Deficient Cancers: The Role of Mutations That Impair Iron Sulfur Cluster Delivery. J Natl Cancer Inst 108 (2016).

40. Q. Guo et al., Succinate dehydrogenase-deficient cancer cells have increased susceptibility to Ym155-induced DNA damage. Endocr Relat Cancer 33 (2026).

41. K. Engle, G. Kumar, Cancer multidrug-resistance reversal by ABCB1 inhibition: A recent update. Eur J Med Chem 239, 114542 (2022).

42. S. S. Thorgeirsson et al., Expression of the multidrug-resistant gene in hepatocarcinogenesis and regenerating rat liver. Science 236, 1120–1122 (1987).

43. M. M. Gottesman, I. Pastan, Biochemistry of multidrug resistance mediated by the multidrug transporter. Annu Rev Biochem 62, 385–427 (1993).

44. A. Sajid, H. Rahman, S. V. Ambudkar, Advances in the structure, mechanism and targeting of chemoresistance-linked ABC transporters. Nat Rev Cancer 23, 762–779 (2023).

45. A. Sajid et al., Reversing the direction of drug transport mediated by the human multidrug transporter P-glycoprotein. Proc Natl Acad Sci U S A 117, 29609–29617 (2020).

46. K. Yasui et al., Alteration in copy numbers of genes as a mechanism for acquired drug resistance. Cancer Res 64, 1403–1410 (2004).

47. R. W. Robey et al., Revisiting the role of ABC transporters in multidrug-resistant cancer. Nat Rev Cancer 18, 452–464 (2018).

48. L. Homolya, M. Hollo, M. Muller, E. B. Mechetner, B. Sarkadi, A new method for a quantitative assessment of P-glycoprotein-related multidrug resistance in tumour cells. Br J Cancer 73, 849–855 (1996).

49. S. V. Ambudkar et al., Biochemical, cellular, and pharmacological aspects of the multidrug transporter. Annu Rev Pharmacol Toxicol 39, 361–398 (1999).

50. Q. Guo et al., TMEM127 suppresses tumor development by promoting RET ubiquitination, positioning, and degradation. Cell Rep 42, 113070 (2023).

51. A. Flynn et al., Multi-omic analysis of SDHB-deficient pheochromocytomas and paragangliomas identifies metastasis and treatment-related molecular profiles. Nat Commun 16, 2632 (2025).

52. P. Kantharidis et al., Altered methylation of the human MDR1 promoter is associated with acquired multidrug resistance. Clin Cancer Res 3, 2025–2032 (1997).

53. A. G. Manjon et al., Perturbations in 3D genome organization can promote acquired drug resistance. Cell Rep 42, 113124 (2023).

54. L. Desiderato, M. W. Davey, A. A. Piper, Demethylation of the human MDR1 5’ region accompanies activation of P-glycoprotein expression in a HL60 multidrug resistant subline. Somat Cell Mol Genet 23, 391–400 (1997).

55. A. Li et al., Hypermethylation of ATP-binding cassette B1 (ABCB1) multidrug resistance 1 (MDR1) is associated with cisplatin resistance in the A549 lung adenocarcinoma cell line. Int J Exp Pathol 97, 412–421 (2016).

56. K. Reed et al., Hypermethylation of the ABCB1 downstream gene promoter accompanies ABCB1 gene amplification and increased expression in docetaxel-resistant MCF-7 breast tumor cells. Epigenetics 3, 270–280 (2008).

57. M. S. Williams, F. M. Amaral, F. Simeoni, T. C. Somervaille, A stress-responsive enhancer induces dynamic drug resistance in acute myeloid leukemia. J Clin Invest 130, 1217–1232 (2020).

58. Y. G. Park et al., Outcomes of systemic treatment according to germline mutational status in patients with metastatic pheochromocytoma and paraganglioma. Clin Genitourin Cancer 22, 413–419 (2024).

59. C. G. Duncan et al., A heterozygous IDH1R132H/WT mutation induces genome-wide alterations in DNA methylation. Genome Res 22, 2339–2355 (2012).

60. X. Zhang et al., Comprehensive molecular profiling of FH-deficient renal cell carcinoma identifies molecular subtypes and potential therapeutic targets. Nat Commun 16, 4398 (2025).

61. W. Xu et al., Oncometabolite 2-hydroxyglutarate is a competitive inhibitor of alpha-ketoglutarate-dependent dioxygenases. Cancer Cell 19, 17–30 (2011).

62. A. Majchrzak-Celinska, A. Sidhu, I. Miechowicz, W. Nowak, A. M. Barciszewska, ABCB1 Is Frequently Methylated in Higher-Grade Gliomas and May Serve as a Diagnostic Biomarker of More Aggressive Tumors. J Clin Med 11 (2022).

63. C. J. Ricketts et al., Kidney tumors associated with germline mutations of FH and SDHB show a CpG island methylator phenotype (CIMP). PLoS One 17, e0278108 (2022).

64. A. Yagoda, D. Petrylak, S. Thompson, Cytotoxic chemotherapy for advanced renal cell carcinoma. Urol Clin North Am 20, 303–321 (1993).

65. D. E. Johnson, L. Rodriguez, P. Y. Holoye, M. L. Samuels, Combination vincristine (NSC-67574) and hydroxyurea (NSC-32065) for metastatic renal carcinoma. Cancer Chemother Rep 59, 1159–1160 (1975).

66. B. van der Werf-Messing, J. Mulder, Letter: Metastatic kidney cancer treated with multiple drug therapy at the Rotterdam Radiotherapy Institute. Br J Cancer 29, 491–492 (1974).

67. C. Annunziata et al., A versatile method for the identification of senolytic compounds. Cell Stress 7, 105–111 (2023).

68. E. S. Potekhina et al., Drug Screening with Genetically Encoded Fluorescent Sensors: Today and Tomorrow. Int J Mol Sci 22 (2020).

69. H. Enokida et al., CpG hypermethylation of MDR1 gene contributes to the pathogenesis and progression of human prostate cancer. Cancer Res 64, 5956–5962 (2004).

70. V. Indio et al., Gene Expression Landscape of SDH-Deficient Gastrointestinal Stromal Tumors. J Clin Med 10 (2021).

71. A. Flynn et al., Multi-omic analysis of SDHB-deficient pheochromocytomas and paragangliomas identifies metastasis and treatment-related molecular profiles. Res Sq 10.21203/rs.3.rs-4410500/v1 (2024).

72. M. Sciacovelli et al., Fumarate is an epigenetic modifier that elicits epithelial-to-mesenchymal transition. Nature 537, 544–547 (2016).

